# Bromodomain inhibition reveals FGF15/19 as a target of epigenetic regulation and metabolic control

**DOI:** 10.1101/2019.12.11.872887

**Authors:** Chisayo Kozuka, Vicencia Sales, Soravis Osataphan, Yixing Yuchi, Jeremy Chimene-Weiss, Christopher Mulla, Elvira Isganaitis, Jessica Desmond, Suzuka Sanechika, Joji Kusuyama, Laurie Goodyear, Xu Shi, Robert E. Gerszten, Lei Wu, Jun Qi, Mary-Elizabeth Patti

## Abstract

Epigenetic regulation is an important factor in glucose metabolism, but underlying mechanisms remain largely unknown. Here we demonstrated that bromodomain-containing proteins (Brds), transcriptional regulators binding to acetylated histone, are potent modulators of glucose metabolism via the gut-liver farnesoid X receptor (FXR)-fibroblast growth factor 15/19 (FGF15/19) pathway. In vivo inhibition of Brd4 by the inhibitor JQ1 in mice strongly inhibited ileal expression of FGF15, resulting in decreased FGFR4-related signaling, increased glucose production in the liver and hyperglycemia. Adverse metabolic effects of BRD4 inhibition were reversed by overexpression of FGF19, with improvement in hyperglycemia. At a cellular level, we demonstrate that BRD4 binds to the promoter region of FGF19 in human intestinal cells; BRD inhibition by JQ1 reduces binding to the FGF19 promoter and downregulates FGF19 expression. Thus, we identify Brd4 as a novel transcriptional regulator of intestinal FGF15/19 in ileum, and a contributor to hepatic and systemic glucose metabolism.

## Introduction

Type 2 diabetes (T2D) is a complex disorder influenced by interactions between multiple genetic loci and environmental factors (Pinney and Simmons, 2010). Environmental factors contributing to metabolic disease, such as intrauterine environment, diet and physical activity, may mediate risk via epigenetic mechanisms, such as DNA methylation, histone modification, and noncoding RNAs. Given that epigenetic mediators are influenced by metabolic signals and in turn modulate transcriptional and/or developmental responses, understanding mechanisms by which epigenetic signals influence metabolic disease risk is a key scientific challenge.

One key epigenetic mark is histone acetylation, which mediates chromatin accessibility to transcriptional factors and coactivators (Verdin and Ott, 2015). In turn, histone acetylation is mediated by histone acetyltransferase (HAT) and removed by histone deacetylase (HDAC). Modulation of histone acetylation can alter systemic metabolism. For example, mice with heterozygous deficiency of the HAT CREB-binding protein (CBP), remain insulin sensitive, despite lipodystrophy (Yamauchi et al., 2002). Conversely, deletion of HDAC in skeletal muscle increases lipid oxidation, energy expenditure and insulin resistance (Gaur et al., 2016; Hong et al., 2017). Together, these data demonstrate that histone acetylation plays an important role in energy expenditure and glucose metabolism.

The Bromodomain and Extra-Terminal Domain (BET) family of proteins has two tandem bromodomains that recognize acetylated lysine of histones or non-histone targets. The mammalian BET family comprises bromodomain-containing proteins 2 (Brd2), Brd3, Brd4 and BrdT in mammals, which are “readers” that bind to acetylated histones and recruit transcription factors (Marmorstein and Zhou, 2014). While Brds are recognized for their potential as a target in cancer therapeutics, Brds also influence metabolism. For example, mice with genetic disruption of Brd2 have severe obesity, but normal glucose metabolism, potentially via increased peroxisome-proliferator-activated receptor (PPAR)-γ activity (Wang et al., 2009). Brd4 binds to enhancers regulating adipogenesis and myogenesis (Lee et al., 2017). Moreover, Brd4 is a coactivator of nuclear factor-κB, (Huang et al., 2009; Mauro et al., 2011), and inhibition of Brd4 can modulate OXPHOS capacity (Barrow et al., 2016). However, the role of BET proteins in systemic metabolism remains ill-defined.

Mouse fibroblast growth factor (FGF) 15 and its human ortholog FGF19 share about 50% amino acid identity and have similar physiological functions to regulate intestine-to-liver crosstalk via a complex feedback loop (Markan and Potthoff, 2016). Bile acids are synthesized from cholesterol in the liver and enter the enterohepatic circulation. After entry into the intestinal lumen, bile acids are transported into enterocytes by apical sodium dependent bile acid transporter (ASBT) and activate the farnesoid X receptor (FXR) to promote FGF15/19 gene transcription and secretion of FGF15/19 into the circulation. In the liver, FGF15/19 binds to and activates the FGF receptor 4 (FGFR4)/β-klotho receptor complex, leading to suppression of bile acid synthesis via repression of the rate-limiting enzyme cytochrome P450 7A1 (CYP7A1). FGF15/19 also exerts potent effects on glucose metabolism, reducing blood glucose via increased activity of Agouti-related peptide/Neuropeptide Y neurons in hypothalamus (Liu et al., 2018; Morton et al., 2013) and modulation of hepatic metabolism, including inhibition of gluconeogenesis and lipogenesis and increased glycogen synthesis (Owen et al., 2015). We now demonstrate that the FGF15/19 signaling pathway is a target of epigenetic modification by the bromodomain inhibitor, JQ-1.

## Results

### Brd4 inhibition by JQ-1 decreases body weight and induces hyperglycemia

JQ-1, a Brd inhibitor, shows high selectivity for Brd4 (Filippakopoulos et al., 2010). To determine the effect of Brd4 inhibition on body weight and glucose metabolism, we treated CD1 mice with JQ-1 (25 mg/kg intraperitoneally) for 11 days (Figure 1A). This dose was previously shown to be without toxic effects during long term treatment (Filippakopoulos et al., 2010). JQ-1 decreased body weight modestly (Veh: 41.4 ± 0.9 g; JQ-1: 36.8 ± 0.5 g, *p*<0.01) (Figure 1B). Modest effects on food intake were observed in one cohort (Veh: 5.4 ± 0.2 g/day; JQ-1: 3.7 ± 1.1 g/day), but not in subsequent cohorts of mice treated with the same dose of JQ-1. Blood glucose was significantly increased in JQ-1 treated mice vs. vehicle, both after a 4-hour fast (Veh: 186 ± 17 mg/dl; JQ-1: 264 ± 14 mg/dl, *p*<0.01, Figure 1C) and during intraperitoneal glucose tolerance testing (peak glucose: Veh: 384 ± 18 mg/dl; JQ-1: 592 ± 8 mg/dl, *p*<0.01) (Figure 1D-E).

**Figure 1.**
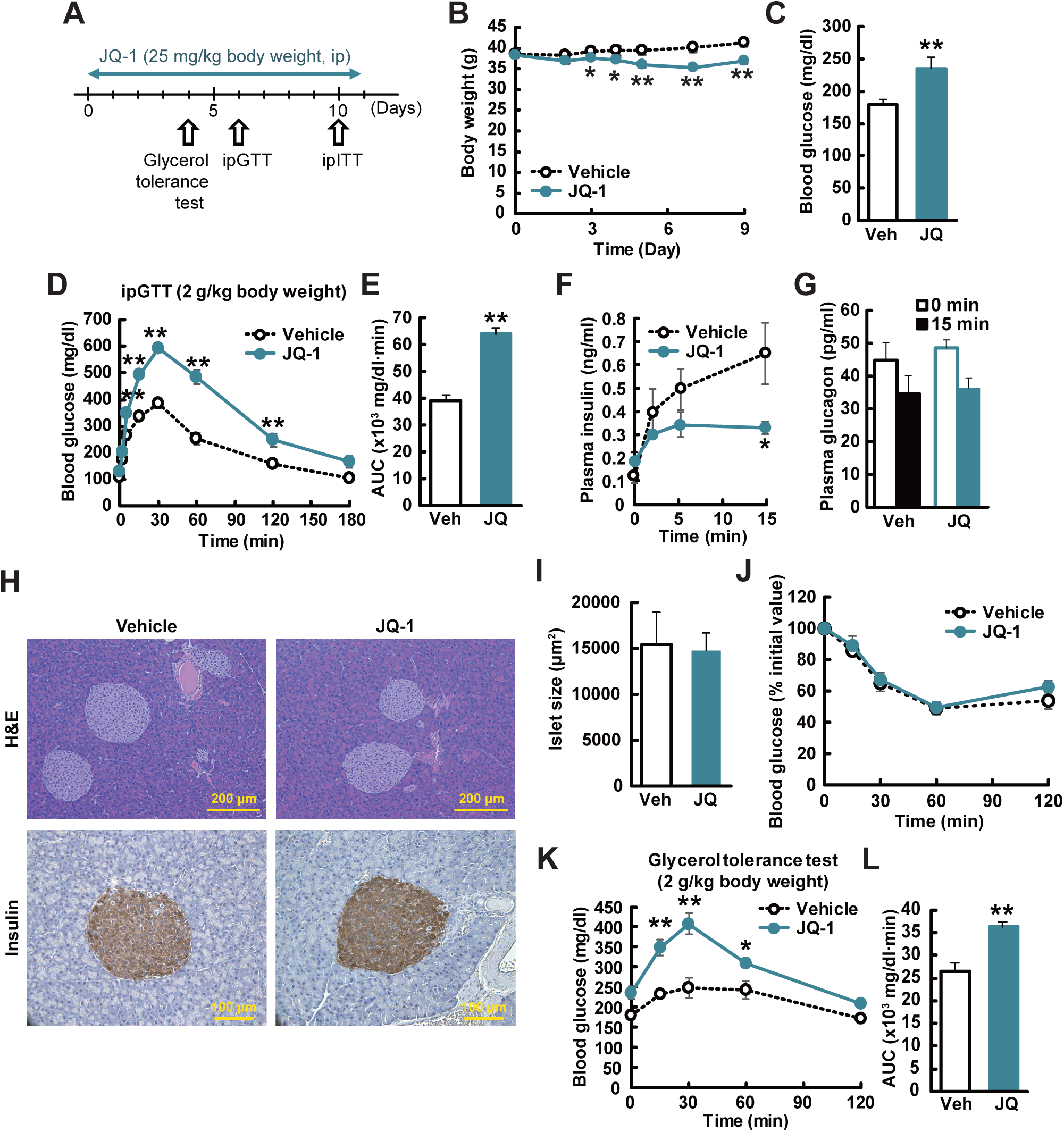
JQ-1 induces hyperglycemia *in vivo*. (**A**) Experimental protocol. (**B**) Body weight changes during JQ-1 treatment (*n* = 5-6). (**C**) Fasting blood glucose levels (4-h fasting, day 10, *n* = 5-6). (**D, E**) Blood glucose levels and area under the curve (AUC) during ipGTT (day 6). (**F, G**) Plasma insulin and glucagon levels during ipGTT. (**H**) H&E and insulin staining of pancreatic sections. Scale bar, 200 and 100 μm. (**I**) Islet size (*n* = 5-6; >100 islets/mouse). (**J, K**) Intraperitoneal insulin tolerance test (day10) and glycerol tolerance test (day 4). \**P* < 0.05, \*\**P* < 0.01, *vs.* vehicle-treated mice (Veh). Data are expressed as means ± SEM.

We next analyzed potential contributors to hyperglycemia in JQ-1-treated mice. While fasting plasma insulin did not differ, insulin levels were significantly decreased in JQ-1-treated mice after glucose injection (50% reduction at 15 minutes, p<0.05) (Figure 1F). Glucose similarly reduced glucagon levels in both vehicle and JQ-1-treated mice (Figure 1G). JQ-1 had no effect on pancreatic islet size (Veh; 15,427 ± 3,503 μm^2^, JQ; 14,763 ± 1,928 μm^2^) (Figure 1H, I). There was no difference in insulin sensitivity as determined by ITT (Figure 1J). Glycerol tolerance testing revealed greater increase in blood glucose in JQ-1-treated mice vs. vehicle-treated mice (Figure 1K, L).

Fasting hyperglycemia in JQ-1-treated mice without change in fasting insulin and increased glycemic response to glycerol suggested increased gluconeogenesis. We therefore asked whether this was related to impaired insulin action or cell autonomous insulin resistance. Hepatic insulin action in vivo, as measured by Akt phosphorylation (**Figure S1A-C**) did not differ. Moreover, basal glucose production in primary hepatocytes did not differ with JQ-1, and the effect of insulin to suppress glucose production was only modestly reduced (vehicle vs. JQ-1, *p* = 0.05, **Figure S1D**). Together, these data suggested that reduction in systemic insulin sensitivity or cell autonomous hepatic insulin action was not likely the dominant mediator of in vivo hyperglycemia.

### JQ-1 alters hepatic gene expression and sterol metabolism

To identify potential molecular mediators of fasting hyperglycemia *in vivo*, we analyzed gene expression in the liver using microarray. 1471 genes were upregulated and 1036 genes were downregulated by JQ-1 (FDR<0.25). Pathway analysis using Gene Set Enrichment Analysis (GSEA) and REACTOME pathways (**Table S1**) revealed upregulation of multiple pathways related to glucose production and lipid metabolism (biosynthesis of triglycerides, fatty acids, and cholesterol; β-oxidation) in JQ-1-treated mice (**Figure S1E, F**). Analysis of individual genes revealed increased expression of genes related to gluconeogenesis (fold change vs. vehicle: *Pepck*: 1.6 ± 0.2, *p* < 0.01, *G6pase*: 2.0 ± 0.6, **Figure S1G, H**). Effects on lipid metabolism were more striking, with significant increases in expression of genes regulating FA synthesis (*Acc*: 1.9 ± 0.2, *p* < 0.01; *Fasn* 2.2 ± 0.3, *p* < 0.01), β-oxidation (*Ppar*α: 1.6 ± 0.1, *p* < 0.01; *Cpt1*α: 1.2 ± 0.0, *p* < 0.01), and cholesterol synthesis (e.g. *Hmgcr*, *Hmgcs1*, and *Srebp2*) (**Figure S1I-O**).

Given these prominent patterns of lipid-related gene expression in JQ-1 treated mice, we analyzed plasma and hepatic lipids. There were no differences in either plasma or hepatic triglycerides (**Figure S2A, B**). By contrast, cholesterol levels were reduced by 52% in plasma (p<0.05), with 28% reduction in liver content in JQ-1-treated mice (plasma: veh; 165 ± mg/dl, JQ-1; 79 ± 29 mg/dl, *p* < 0.05; liver: veh; 0.94 ± 0.1 mg/g tissue, JQ-1; 0.68 ± 0.1 mg/g tissue, *p* < 0.05) (**Figure S2C, D**).

Reductions in both plasma and tissue cholesterol levels, despite significantly increased expression of cholesterol synthetic genes, suggested that cholesterol catabolic pathways, such as bile acid metabolism, may be upregulated. Indeed the major enzymes of both classic and alternative bile acid synthesis pathways were upregulated by JQ-1, with a 1.4-fold increase in the rate-limiting enzyme *Cyp7a1* (*p* = 0.08) and 1.2-fold increase in *Cyp27a1* (*p* < 0.01) (**Figure S1P, Q and Figure S2E**). However, there was no net difference in hepatic and plasma bile acid composition between vehicle and JQ-1-treated mice (**Figure S2F, G**), potentially linked to downregulation of the downstream enzymes Cyp7b1 and Cyp8b1 (reduced by 56 and 46% respectively, **Figure S2E**).

The complex pattern of lipid, sterol, and bile acid gene expression and metabolism in JQ-1-treated liver raised the possibility that FGF receptor-dependent signaling might be reduced by JQ-1. Indeed, 10 of 14 top-ranking pathways *downregulated* in JQ-1-treated mice were related to FGFR signaling (FDR <0.25, Figure 2A). Expression of Fgfr4 as determined by PCR was decreased by 27% (*p* < 0.01) with a similar trend for its coreceptor beta-Klotho (17% lower, *p* = 0.07). Moreover, Fgfr signaling was significantly reduced with a 46% reduction in phosphorylation of FGF receptor substrate 2 (FRS2) in liver protein extracts of JQ-1 treated mice (*p* < 0.01, **Figure S1A, B**).

**Figure 2.**
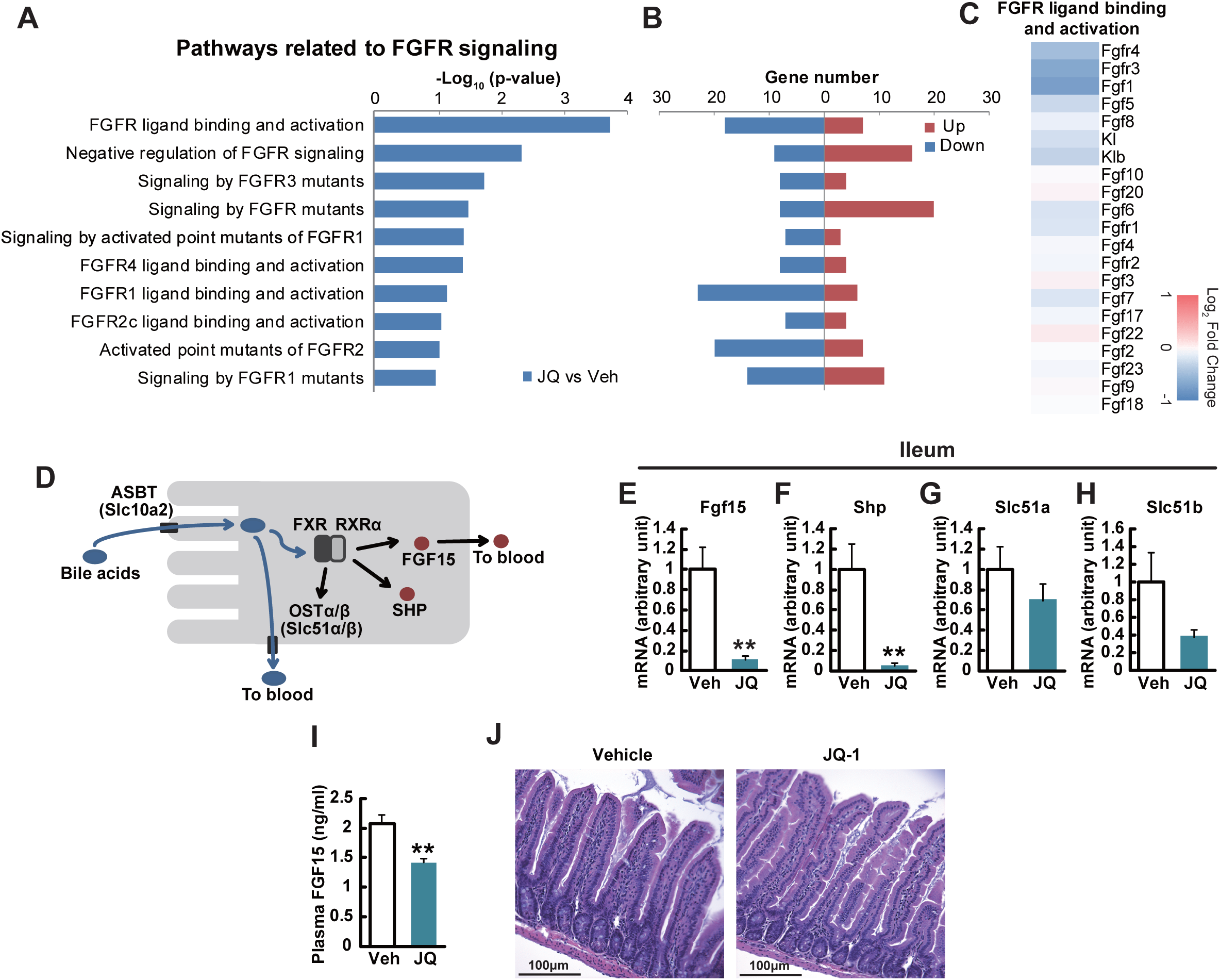
JQ-1 modulates hepatic transcription related to Fgfr signaling, reduces Fgfr signaling, and reduces ileal Fgf15 and Shp expression in vivo. (**A-C**) Microarray analysis in the liver was performed in JQ-1 vs. vehicle-treated mice (*n* = 3 per group). Minus log_10_ (p-value) (**A**) and upregulated or downregulated gene numbers (**B**) in pathways related to FGFR signaling. (**C**) Heatmap demonstrate log_2_ fold change (JQ-1 vs. vehicle) in gene expression in pathways related to FGFR ligand binding and activation. (**D**) Schematic of bile acid signaling in ileal enterocytes. (**E-H**) Expression levels of genes related to bile acid signaling in ileum (*Fgf15* (**E**), *Nr0b2* (SHP) (**F**), *Slc51a* (OSTα) (**G**) and *Slc51b* (OSTβ) (**H**)) (*n* = 5-6). Expression levels were normalized by those of *Rpl13*. (**I**) Plasma FGF15 level in JQ-1 vs. vehicle-treated mice (*n* = 5-6). \**P* < 0.05, \*\**P* < 0.01, *vs.* vehicle-treated mice (Veh). Data are expressed as means ± SEM. (**J**) H&E staining of intestinal sections. Scale bar, 100 μm.

### Brd4 inhibition by JQ-1 reduces ileal expression of Fgf15

Upregulation of bile acid synthetic enzymes and downregulation of FGFR-mediated signaling suggest that JQ1 modulated systemic and hepatic lipid and bile acid metabolism via the complex regulatory loop involving the bile acid-responsive hormone FGF15/19. FGF15 (FGF19 in humans) is produced in intestinal enterocytes in response to bile acids, secreted into the bloodstream, and binds FGF receptors in the liver to inhibit both bile acid synthesis and cholesterol metabolism (Kim et al., 2015; Kliewer and Mangelsdorf, 2015).

Luminal bile acids are transported into enterocytes by ASBT (Slc10a2), where they can bind to FXR and activate its transcriptional activity to increase expression of its targets including *Fgf15*, *Shp*, *Slc15a* and *Slc51b* (Wong et al., 2011). While there was no change in no change in Slc10a2, Tgr5, Fxr, or RXRa expression in ileum of JQ-1-treated mice (not shown), we observed profound alterations in expression of FXR target genes. Expression of *Fgf15* and *Shp* was reduced by 90% and 95% (*p*<0.01 for all), with similar trend for *Slc51a* and *Slc51b* (Figure 2E-H). In agreement, plasma FGF15 was decreased by 29% in JQ-1-treated mice (veh; 2.1 ± 0.1 ng/ml, JQ-1; 1.4 ± 0.1 ng/ml, *p* < 0.05) (Figure 2I).

We next evaluated potential mechanisms mediating JQ-1 decreases in Fgf15 expression. Previous studies have shown that RNAi-mediated silencing of Brd4 reduces intestinal cellular diversity (Bolden et al., 2014), potentially via reduced intestinal stem cell differentiation (Nakagawa et al., 2016); thus, alterations in number or function of specific intestinal cell subtypes could potentially contribute to JQ-1-mediated changes in intestinal FGF15 levels. Histologic analysis revealed no change in the overall villus structure in JQ-1-treated mice (Figure 2J). However, JQ-1 was associated with reduced numbers of Paneth cells, as previously reported (Bolden et al., 2014) (**Figure S3A**). In parallel, mRNA levels of Reg4, a Paneth cell marker, were decreased by JQ-1, but expression of the microbicidal functional marker α-defensin cryptdin-4 (Crp4) did not differ (**Figure S3G, K**) (Ouellette, 2011). Moreover, there was no change in expression of other cell type markers for enterocytes (keratin 20 (Krt20)), enteroendocrine cells (hepatocyte nuclear factor (HNF) 1α), neuroendocrine cells (chromogranin A (ChgA)), goblet cells (mucin 2 (Muc2)), stem cells (leucine-rich orphan G-protein-coupled receptor 5 (Lgr5)), or intestinal neuroendocrine peptides (*Gcg* or *Pyy*), suggesting JQ-1-mediated reduction in Fgf15 expression was not related to global alterations in cell type distribution (**Figure S3B-J**).

### JQ-1 inhibits Brd4 binding to the Fgf15 promoter in HT-29 cells

Marked downregulation of ileal expression of Fgf15 in JQ-1 treated mice suggested that the bile acid-Fgf15 feedback loop might play a prominent role in systemic metabolic effects of JQ-1. To test this hypothesis at a cellular level, we utilized the human intestinal cell line HT-29, which constitutively expresses FGF19 at a high level (Vergnes et al., 2013). HT-29 cells were treated with 500 nM JQ-1 for 24 hours. While there was no change in *BRD4* expression (Figure 3A), expression of the Brd4 target gene *MYC* was reduced by 83% with JQ-1 (*p* < 0.01), as predicted (Figure 3B). In parallel, expression of *FGF19* was reduced by 96% (*p* < 0.01) with similar dramatic reduction in *SHP* (89% reduction, *p* < 0.01) (Figure 3C, D). There was no change in *FXR* or *SLC51B* expression (data not shown), consistent with prior evidence indicating BRD4 does not bind to FXR or SLC51B (GSE73319 (McCleland et al., 2016) and ENCSR514EOE (Consortium, 2012). Moreover, secretion of FGF19 into culture medium was completely abolished by JQ-1, with >98% decrease in the conditioned medium (*p*<0.01, Figure 3E). To determine whether JQ-1 mediated reduction in expression of FGF19 and SHP was mediated by modulation of BRD4 binding to promoter regions of these genes, we performed ChIP-PCR analysis using an anti-BRD4 antibody (Figure 3F) (Rathert et al., 2015). As expected, BRD4 bound to its target Myc (control); Brd4 also bound robustly to promoter sequences of both FGF19 and SHP (Figure 3G, H). This was markedly inhibited by JQ-1, with a >90% reduction in binding (Figure 3G, H).

**Figure 3.**
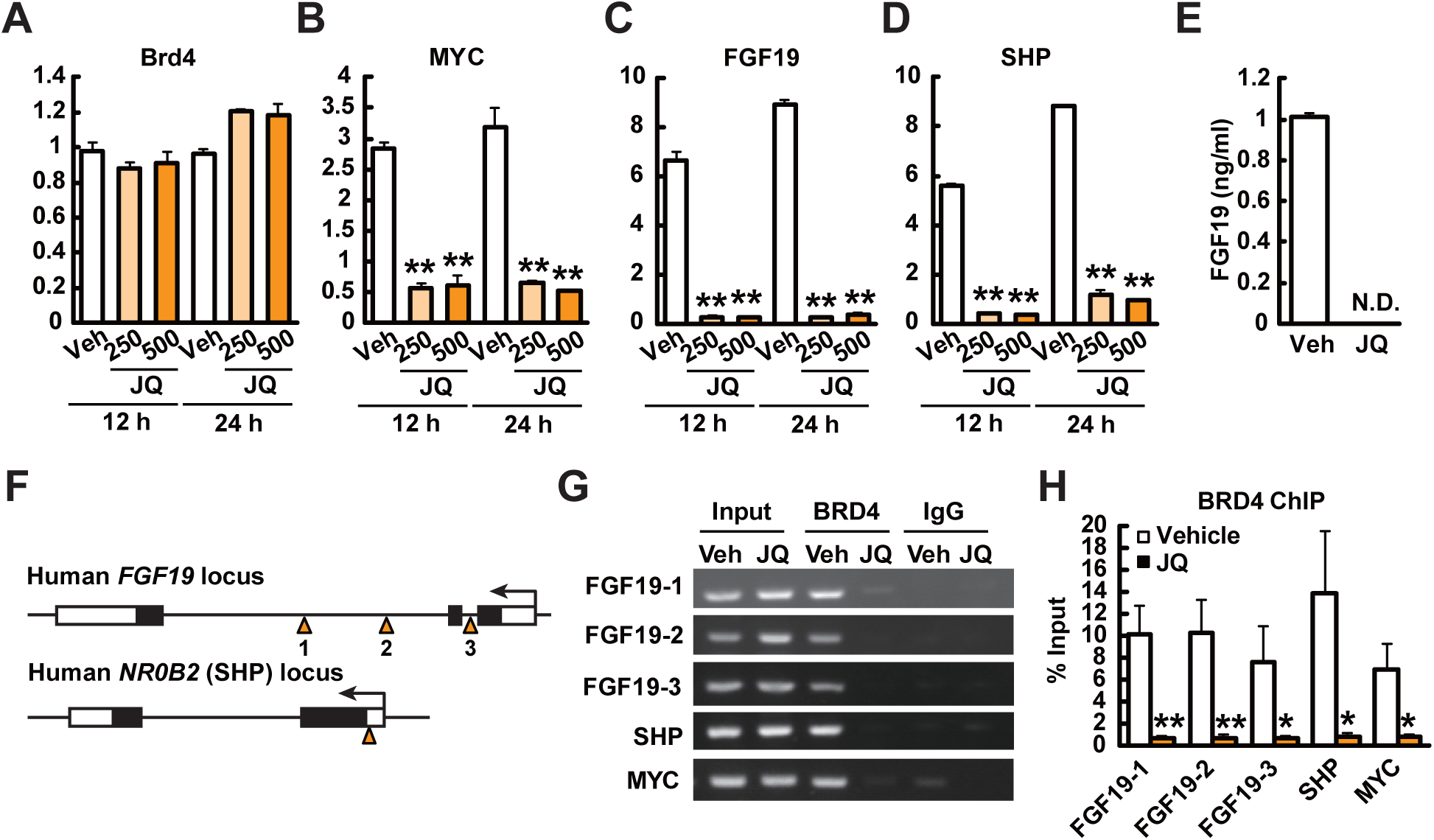
BRD4 binds promoter regions of FGF19 and SHP to modulate expression in HT-29 cells. HT-29 cells were treated with 250 or 500 nM JQ-1 or DMSO (Veh) for 12-24 h. (**A-D**) Gene expression levels of *BRD4* (**A**), *MYC* (**B**), *FGF19* (**C**), and *NR0B2* (SHP) (**D**) (*n* = 4). The levels were normalized by those of *RN18S*. (**E**) FGF19 level in conditioned media (*n* = 4). (**F**) Primer design for ChIP assays for *FGF19* and *NR0B2* (SHP) promoter regions. (**G-H**) Cells were incubated with JQ-1 or DMSO (Veh) for 24 h and then harvested for the ChIP assay. The precipitated DNA was analyzed by PCR (**G**) and real-time PCR (**H**) (*n* = 4). \**P* < 0.05, \*\**P* < 0.01, *vs.* vehicle-treated mice (Veh). Data are expressed as means ± SEM.

### Overexpression of FGF19 reverses glucose intolerance induced by JQ-1

To determine the requirement for FGF15/19 signaling in mediating the metabolic effects of JQ-1, we treated mice with AAV-FGF19 or GFP control vectors, and then treated mice with JQ-1 (25 mg/kg, ip) for 10 days (Figure 4A). Plasma FGF19 was not detectable in AAV-GFP controls, but was readily detected in plasma of AAV-FGF19 mice (Figure 4B). Consistent with prior studies (Lan et al., 2017; Morton et al., 2013), AAV-FGF19-treated mice had lower body weight and blood glucose levels at baseline (Figure 4C) but weight did not change during JQ-1 treatment in either GFP or FGF19 groups (Figure 4C). As with prior cohorts, JQ-1 increased blood glucose in AAV-GFP control mice as early as 4 days after the onset of treatment, but this effect was not observed in AAV-FGF19 mice. Glucose tolerance testing at day 10 of treatment again revealed significant impairment in JQ-1 treated mice (35% increase in glucose AUC, *p* < 0.01) (Figure 4D). By contrast, the ability of JQ-1 to impair glucose tolerance was fully reversed in mice overexpressing FGF19 (Figure 4D, E). Moreover, overexpression of FGF19 reduced both basal and glucose-stimulated insulin levels (Figure 4F), indicating FGF19-mediated reversal of glucose tolerance in JQ-1 treated mice is insulin-independent.

**Figure 4.**
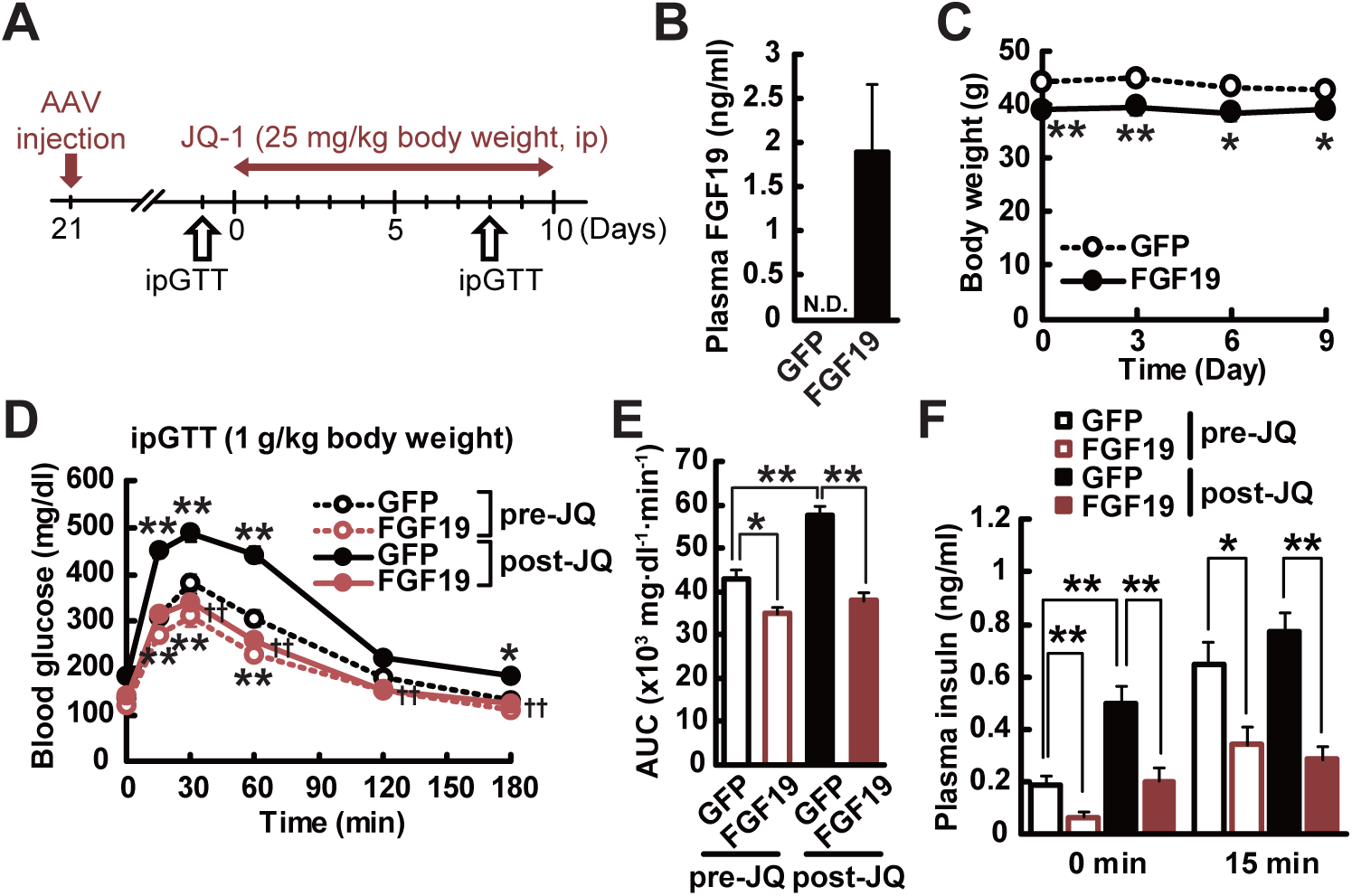
Overexpression of FGF19 reverses hyperglycemia induced by JQ-1. (**A**) Experimental protocol. (**B**) Plasma FGF19 levels in AAV-GFP- and AAV-FGF19-treated mice. (**C**) Body weight in AAV-GFP- and AAV-FGF19-treated mice during JQ-1 treatment. (**D-E**) Blood glucose levels (**D**) and area under the curve (AUC) (**E**) during ipGTT (day10). (**F**) Plasma insulin levels during ipGTT. \**P* < 0.05, \*\**P* < 0.01, *vs.* AAV-GFP-treated mice (GFP). Data are expressed as means ± SEM.

## Discussion

Our studies identify the intestinal Fgf15/19 hormonal axis as a target of epigenetic and transcriptional regulation by the bromodomain protein Brd4 and its inhibitor JQ-1. We demonstrate that pharmacological inhibition of Brd4 induces hyperglycemia and impaired glucose tolerance in mice, via inhibition of intestinal FGF15 expression and reduced Fgfr4-mediated signaling in liver. ChIP-PCR analysis revealed that FGF19 and SHP are novel targets of Brd4 in intestine, and that JQ-1 reduced Brd4 binding to the promoter of these targets, reduced their expression, and reduced secretion of FGF19. Experimental overexpression of FGF19 normalized JQ-1-induced hyperglycemia. Thus, Brd4-dependent transcription is a potent regulator of intestinal endocrine function and systemic glucose metabolism via FGF15/19-dependent mechanisms.

Our data provide new evidence supporting epigenetic regulation of intestinal function and metabolism and identification of new gene targets of Brd4 regulation. Prior studies have demonstrated that genetic or pharmacologic inhibition of Brd4-dependent signaling (Bolden et al., 2014) (Nakagawa et al., 2016) can modulate intestinal cell populations. While we also find that JQ-1 decreased the number of Reg4+ Paneth cells, there was no change in the Paneth cell functional marker Crp4 (Ouellette, 2011) (**Figure S3**) nor in ileal expression of cytokines (data not shown). Moreover, JQ-1 did not alter markers of additional intestinal cell populations. Thus, effects of JQ-1 in our experimental conditions do not appear to require alterations in Paneth cell-linked host defense systems or cell type distribution. Rather, our data point to new targets of Brd4 action in the intestine – including regulation of enterocyte Fgf15/19 expression – with secondary effects on hepatic and systemic glucose and sterol lipid metabolism.

Fgf15/19 expression and secretion in the intestine can be regulated by bile acid binding to the luminal bile acid receptor Tgr5 or the nuclear receptor Fxr. We observed no change in expression of Tgr5, the apical bile acid transporter Slc10a2, Fxr, or Rxr, and plasma levels of bile acids did not differ significantly in JQ-1-treated mice. We cannot exclude the possibility that luminal signaling via bile acids or other metabolites could contribute to reduced intestinal Fgf15 expression and secretion in response to JQ-1 in vivo. However, our data in cultured cells suggest that JQ-1 exerts a direct, cell autonomous effect on transcription and secretion of Fgf15/19 via inhibition of Brd4.

Our data suggest that inhibition of Fgf15/19 is also the dominant mediator of in vivo metabolic effects of JQ-1. Fgf15/19 is increasingly recognized as a potent regulator of systemic glucose metabolism. Direct administration or transgenic overexpression of FGF19 in mice fed either chow or high fat diet improves glucose tolerance, despite lower insulin levels (Fu et al.; Tomlinson et al., 2002), and reverses obesity, hyperglycemia, and insulin resistance in mice with genetic obesity (Fu et al., 2004; Morton et al., 2013). Such improvements in systemic glucose metabolism may result in part from FGF-19 effects to increase insulin-independent glucose uptake (Morton et al.) and reduce adiposity (Tomlinson et al., 2002). Interestingly, increased plasma levels of Fgf19 have been implicated as a mediator of the potent impact of bariatric and intestinal surgery to reduce blood glucose levels, with sustained remission of T2D (Bozadjieva et al., 2018) and development of post-bariatric hypoglycemia in some patients (Mulla et al., 2019).

A key mediator of systemic effects of FGF15/19 in normal physiology is altered liver metabolism, with reduced gluconeogenesis and lipogenesis and increased glycogen synthesis (Nies et al., 2015). Consistent with reduction in hepatic signaling by Fgf15/19 in JQ-1-treated mice, we observed (1) downregulation of Fgfr4 and β-Klotho (Klb) pathways and reduced signaling, as indicated by reduced Frs2 phosphorylation (Figure 2A-F), (2) increased hepatic glucose production, as indicated by increased glucose after glycerol injection (Figure 1K) and increased gluconeogenic gene expression (**Supplemental Figure 1**), (3) reduced plasma and hepatic cholesterol, and (4) global changes in the hepatic transcriptome within both glucose and lipid metabolic pathways. Given that the effects of JQ-1 in primary hepatocytes were small in magnitude, cell autonomous effects in the liver do not appear to be dominant. Rather, JQ-1 mediated repression of intestinal FGF15 expression, secretion, and signaling drives in vivo systemic and hepatic glucose metabolism, as demonstrated by reversal of hyperglycemia with restoration of systemic Fgf19 levels.

We acknowledge that JQ-1 may have additional Fgf15/19-independent effects. Neither systemic insulin tolerance nor hepatic insulin signaling were affected by short-term treatment with JQ-1, indicating insulin resistance was not a major contributor to hyperglycemia despite prior reports of Brd4 effects on adipogenesis and myogenesis (Lee et al., 2017). While there were no differences in pancreatic islet size or insulin staining, glucose-stimulated insulin levels were lower in JQ-1-treated mice. While BRD4 can modulate senescence-associated genes in islets in the setting of autoimmunity in mice (Thompson et al., 2019), our data indicate that effects of JQ-1 on insulin secretion are not dominant in our model, as FGF19 overexpression reduced insulin levels but still reversed JQ-1-mediated hyperglycemia. Additional systemic or extrahepatic metabolic effects of inhibition of Brd4-mediated transcription (Lee et al., 2017; Thompson et al., 2019) could also contribute to hyperglycemia induced by JQ-1.

In summary, inhibition of the bromodomain protein Brd4 by JQ-1 inhibits FGF15 and SHP expression in ileum, resulting in reduced FGFR signaling in liver, altered sterol metabolism, increased gluconeogenesis, and hyperglycemia. Effects of JQ-1 on glucose metabolism were FGF15/19 dependent, as overexpression of FGF19 reversed hyperglycemia induced by JQ-1. Thus, our studies identify FGF15/19 as a hormonal target of epigenetic regulation potently contributing to systemic metabolic control.

## Supporting information

Brief Highlights

## ACKNOWLEDGMENTS

We gratefully acknowledge grant support from NIH DK106193 (to MEP), R01DK101043 (to LG), R01CA142106 and R01HD093540 (to JQ), and DK036836 (Joslin DRC). CK was supported by Sunstar Foundation, and SO was supported by the Prince Mahidol Award Foundation. We also thank NGM for providing adenoviral FGF19 and GFP.

## METHODS

### Animal Care and Studies

Male CD-1 mice obtained from Envigo (South Easton, MA, USA) were housed (3-5 per cage) in at 24 °C under a 12-hour light/dark cycle, with free access to food and water. All animal experiments were approved by the IACUC at Joslin Diabetes Center and conducted in accordance with the NIH Guide for the Care and Use of Laboratory Animals.

### Metabolic parameters

Glucose tolerance was assessed after intraperitoneal injection of glucose (1 or 2 g/kg body weight after a 16 hour fast. Blood glucose was measured via tail vein sampling at the indicated time points. Insulin tolerance was assessed after intraperitoneal injection of human insulin (0.5 units/kg body weight, Lilly) after a 4-hour fast. Plasma insulin and glucagon were measured using an enzyme-linked immunosorbent assay (ELISA) kit (Crystal Chem and R&D, respectively). Plasma triglyceride (TG) and total cholesterol were measured using colorimetric assays (Cayman Chemical and Cell Biolabs).

### In vivo treatment with the bromodomain inhibitor JQ-1

The Brd4 inhibitor JQ1 was synthesized and purified in the laboratory of Dr. Jun Qi (DFCI). For in vivo experiments, a stock solution (50 mg/mL in DMSO) was diluted to a working concentration of 5 mg/mL in 10% hydroxypropyl β-cyclodextrin (Sigma). Mice were injected at a dose of 25 or 50 mg/kg given intraperitoneally. Vehicle controls were given an equal amount of DMSO in 10% hydroxypropyl β-cyclodextrin. For in vitro experiments, JQ1 was dissolved in DMSO and added to cells at indicated concentrations, with equal volume of DMSO as control.

### Adenoviral overexpression of FGF19

Mice received a single tail vein injection of 3 × 10^11^ vector genome adeno-associated virus (AAV)-FGF19 or a control virus encoding green fluorescent protein (GFP; AAV-GFP). After 16 days of treatment, tissues were collected.

### Quantitative real-time PCR

Total RNA was extracted using Trizol reagent (Thermo Fisher Scientific, Inc., Waltham, MA, USA), and cDNA was synthesized using a High-Capacity cDNA Reverse Transcription Kit (Thermo Fisher Scientific) according to manufacturer’s instructions. Quantitative real-time PCR was performed using SYBR Green (Bio-Rad, Hercules, CA, USA). Expression was normalized by *Rn18s* (18S rRNA), Rpl13a or 36B4. Primer sequences are provided in **Table S2**.

### Microarray analysis

Total RNA was isolated from liver tissue from 4 representative mice per group. RNA quality was assessed using Agilent 2100 bioanalyzer (Agilent Technologies, Palo Alto, CA), and samples were processed for microarray analysis (Affymetrix Mouse Gene 2.0 ST, Molecular Phenotyping Core, Joslin).

### Bioinformatic analysis

For liver transcriptomics and metabolomics datasets, principal component analysis (PCA) revealed that the first principal component was an extraneous source of variation, so it was accounted for as a covariate in linear modeling (Leek et al., 2010). Metabolomics/lipidomics sample weights were unbiasedly estimated (Ritchie et al., 2006) and used in linear modelling. Linear modeling differential analysis was done with the R package limma (Ritchie et al., 2015). Nominal p-values were corrected for multiple testing using the false discovery rate (FDR). Transcriptomic pathway and transcription factor prediction analysis was done using the Roast method (Wu et al., 2010) with pathways defined by Reactome (Fabregat et al., 2018).

### Immunohistochemistry (IHC)

Pancreata were dissected and fixed in 4% paraformaldehyde, embedded in paraffin, and sectioned. Sections were stained using H&E or immunostained for lysozyme (1:1000; ab108508, Abcam) and insulin (1:100; #4590, Cell Signaling technology). Islet area was calculated from analysis of more than 100 islets per mouse (Adobe Photoshop).

### Cell culture

The human intestinal cell line (HT-29) cells (American Type Cell Collection, Manassas, VA, USA) were maintained in McCoy’s 5A medium (ThermoFisher Scientific) supplemented with 10% FBS, 1% PenStrep (ThermoFisher Scientific) in a humidified atmosphere with 5% CO_2_ at 37 °C.

### Primary hepatocyte isolation and measurement of glucose production

Primary hepatocytes were isolated from C57BL/6 mice after liver perfusion with collagenase and seeded 1 × 10^5^ cells/ml into collagen coated plates containing DMEM (ThermoFisher Scientific). The media were changed after 4 hours. Primary hepatocyte experiments were performed the day after isolation. Glucose production in primary hepatocytes was measured as preciously described (Matsumoto and Sakai, 2012). Briefly, after 6 h serum starvation with or without JQ-1 and FGF19, cells were cultured in glucose and phenol red-free DMEM with or without insulin (10, 100 nM), JQ-1 (250 nM) and FGF19 (100 ng/ml) for 5 h. The medium was used to determine glucose concentrations with the Glucose (HK) Assay Kit (Sigma).

### Chromatin immunoprecipitation (ChIP)-PCR

ChIP was performed on HT-29 cells cultured in the presence or absence of JQ1 (250 nM, 24 hr). Chromatin pooled from approximately 1 × 10^6^ HT-29 cells was used for each immunoprecipitation. HT-29 cells were fixed directly on the dish with 1% formaldehyde for 10 minutes followed by quenching with 0.125M glycine for 5 minutes. Chromatin was extracted, followed by shearing on a Tekmar Sonic Disruptor (Cincinnati, OH, USA) (3 cycles, 80% Amp and 6 sec pulse, 5 min on/off). The sonicated chromatin was immunoprecipitated with 5 μg of antibody (anti-BRD4, Bethyl #A301-985A). bound to Dynabeads (Invitrogen) as previously described (Anand et al., 2013). Cross-linking was reversed in immunoprecipitate and input chromatin samples prior to purification of genomic DNA. Target and non-target regions of genomic DNA were amplified by PCR or qRT-PCR in both the immunoprecipitates and input samples using SYBR Green chemistry. Enrichment was calculated as a percentage of input DNA for each sample. ChIP-PCR primer sequences are shown in **Table S2**. ChIP-PCR primers for MYC were previously described (Rathert et al., 2015).

### Metabolomics

Tissues harvested after 11 days of treatment were used for analysis of plasma and liver metabolites, using liquid chromatography-mass spectrometry (LC/MS) to determine metabolites (Roberts et al., 2012). Missing data were imputed with half of the minimum intensity of the metabolite, and the imputed data were quantile normalized and log_2_-transformed.

### Statistical analysis

Data are expressed as mean ± SEM. One-way ANOVA and repeated-measures ANOVA followed by multiple comparison tests (Bonferroni/Dunn method) were used where applicable. Student’s *t*-test was used to analyze the differences between two groups. Differences were considered significant at *P* < 0.05.

---

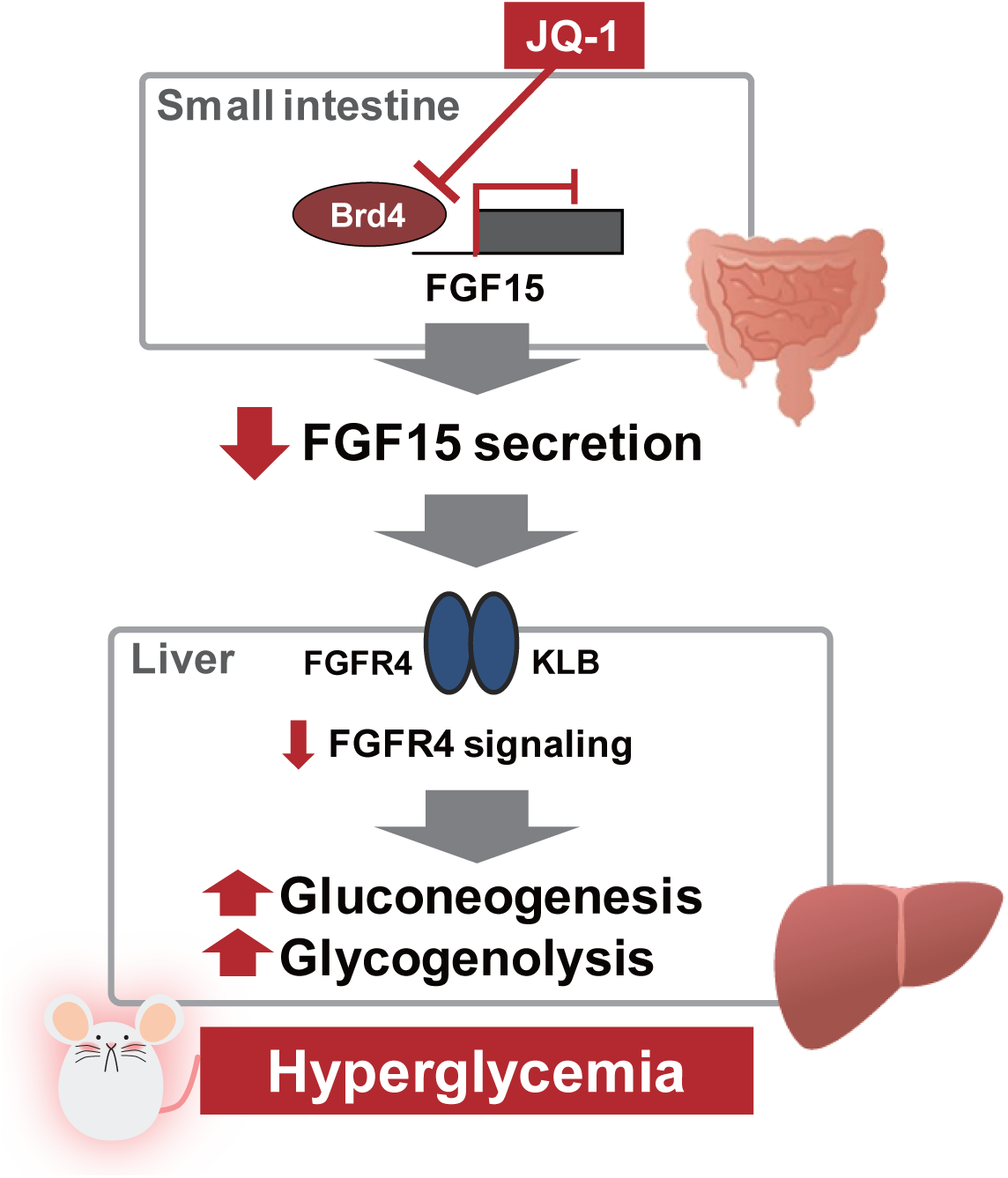

