## Supplemental Tables and Figures for "Bromodomain inhibition reveals FGF15/19 as a target of epigenetic regulation and metabolic control"

**Supplementary Files**

**Supplementary Table 1. Pathway analysis of liver gene expression in JQ1-treated vs. vehicle-treated mice**

**Supplemental Table 2. Primer sequences**

**Mouse**

| **Gene** | **Forward Sequence** | **Reverse Sequence** |
| --- | --- | --- |
| *Rpl13a* | CTGCTCTCAAGGTTGTTCGGCT | CCTTCCGTTTCTCCTCCAGAGT |
| *Slc10a2* (ASBT) | GTCTGTCCCCCAAATGCAACT | CACCCCATAGAAAACATCACCA |
| *Gpbar1* (TGR5) | TGCTTCTTCCTAAGCCTACTACT | CTGATGGTTCCGGCTCCATAG |
| *Nr1h4* (FXR) | CTGGCATCTATGAACTCAGGC | CCATTCGCGGCTTCTTTGG |
| *Rxra* | ATGGACACCAAACATTTCCTGC | CCAGTGGAGAGCCGATTCC |
| *Fgf15* | ACGTCCTTGATGGCAATCG | GAGGACCAAAACGAACGAAATT |
| *Nr0b2* (SHP) | AGGATGCTGTGACCTTCGAG | CAGCTCAAGGCTCCAGAAAGA |
| *Slc51a* (OSTα) | ATGCATCTGGGTGAACAGAA | GAGTAGGGAGGTGAGCAAGC |
| *Slc51b* (OSTβ) | GACCACAGTGCAGAGAAAGC | CTTGTCATCACCACCAGGAC |
| *Pck1* (PEPCK) | CTAACTTGGCCATGATGAACC | CTTCACTGAGGTGCCAGGAG |
| *G6pc* | CTTTGACTCTCTGAAGCCCC | GGGCTAGGCAGTATGGGATA |
| *Acc* | CCTGAAGACCTTAAAGCCAATGC | CCAGCCCACACTGCTTGTA |
| *Fasn* | GCCCGGGTAGCTCTGGGTGTA | TGCTCCCAGCTGCAGGC |
| *Ppara* | CTTCCCAAAGCTCCTTCAAAAA | CTGCGCATGCTCCGTG |
| *Cpt1a* | GCACTGCAGCTCGCACATTACAA | CTCAGACAGTACCTCCTTCAGGAA |
| *Srebp2* | GAACTTTTCCTTAACGTGGGCCT | GAGCATGTCTTCGATGTCGTTCA |
| *Hmgcs1* | AACTGGTGCAGAAATCTCTAGC | GGTTGAATAGCTCAGAACTAGCC |
| *Hmgcr* | AGCTTGCCCGAATTGTATGTG | TCTGTTGTGAACCATGTGACTTC |
| *Nr1h3* (LXR) | AGGAGTGTCGACTTCGCAAA | CTCTTCTTGCCGCTTCAGTTT |
| *Krt20* | CCTGCGAATTGACAATGCTA | CCTTGGAGATCAGCTTCCAC |
| *Hnf1a* | GACCTGACCGAGTTGCCTAAT | CCGGCTCTTTCAGAATGGGT |
| *ChgA* | ATCCTCTCTATCCTGCGACAC | GGGCTCTGGTTCTCAAACACT |
| *Muc2* | AACGATGCCTACACCAAGGTC | ACTGAACTGTATGCCTTCCTCA |
| *Lgr5* | CCTACTCGAAGACTTACCCAGT | GCATTGGGGTGAATGATAGCA |
| *Reg4* | CTGGAATCCCAGGACAAAGAGTG | CTGGAGGCCTCCTCAATGTTTGC |
| *Ctnnb1* (β-catenin) | GCGGCCGCGAGGTACCTGAA | GCAGCTTTTCTGTCCGGCTCCA |
| Gcg | AGGGACCTTTACCAGTGATGT | GCGAATGGCGACTTCTTCTGGGAA |
| Pyy | ACGGTCGCAATGCTGCTAAT | GACATCTCTTTTTCCATACCGCT |
| Crp4 | AAGAGACTAAAACTGAGGAGCAGC | CGGCGGGGGCAGCAGTA |

**Human**

| **Human Gene** | **Forward Sequence** | **Reverse Sequence** |
| --- | --- | --- |
| BRD4 | CATGGACATGAGCACAATCA | TCATGGTCAGGAGGGTTGTA |
| MYC | GGCTCCTGGCAAAAGGTCA | CTGCGTAGTTGTGCTGATGT |
| FGF19 | CGGAGGAAGACTGTGCTTTCG | CTCGGATCGGTACACATTGTAG |
| NR0B2 (SHP) | ATCCTCTTCAACCCCGATGTG | ACTTCACACAGCACCCAGTG |

**ChIP-PCR**

| **ChIP-PCR primers** | **Forward Sequence** | **Reverse Sequence** |
| --- | --- | --- |
| FGF19-1 | GCTGGGACAAAGCATCGGA | TCCCAAAGGACAAGCCAAGTAA |
| FGF19-2 | GTCCCGCTGTTCACAAAGGA | CCACCCCAGAAAGGATGCAG |
| FGF19-3 | GTGCGGTCGGTCCACAA | TTTTGGCACCTGGACGTTAG |
| SHP | TGGCTGGTGCTCATGGTTAG | TAAATAGCACCCACAGCGCA |

**Supplemental Figure 1. JQ-1 modulates expression of genes regulating glucose and lipid metabolism in the liver. *Related to Figure 1***

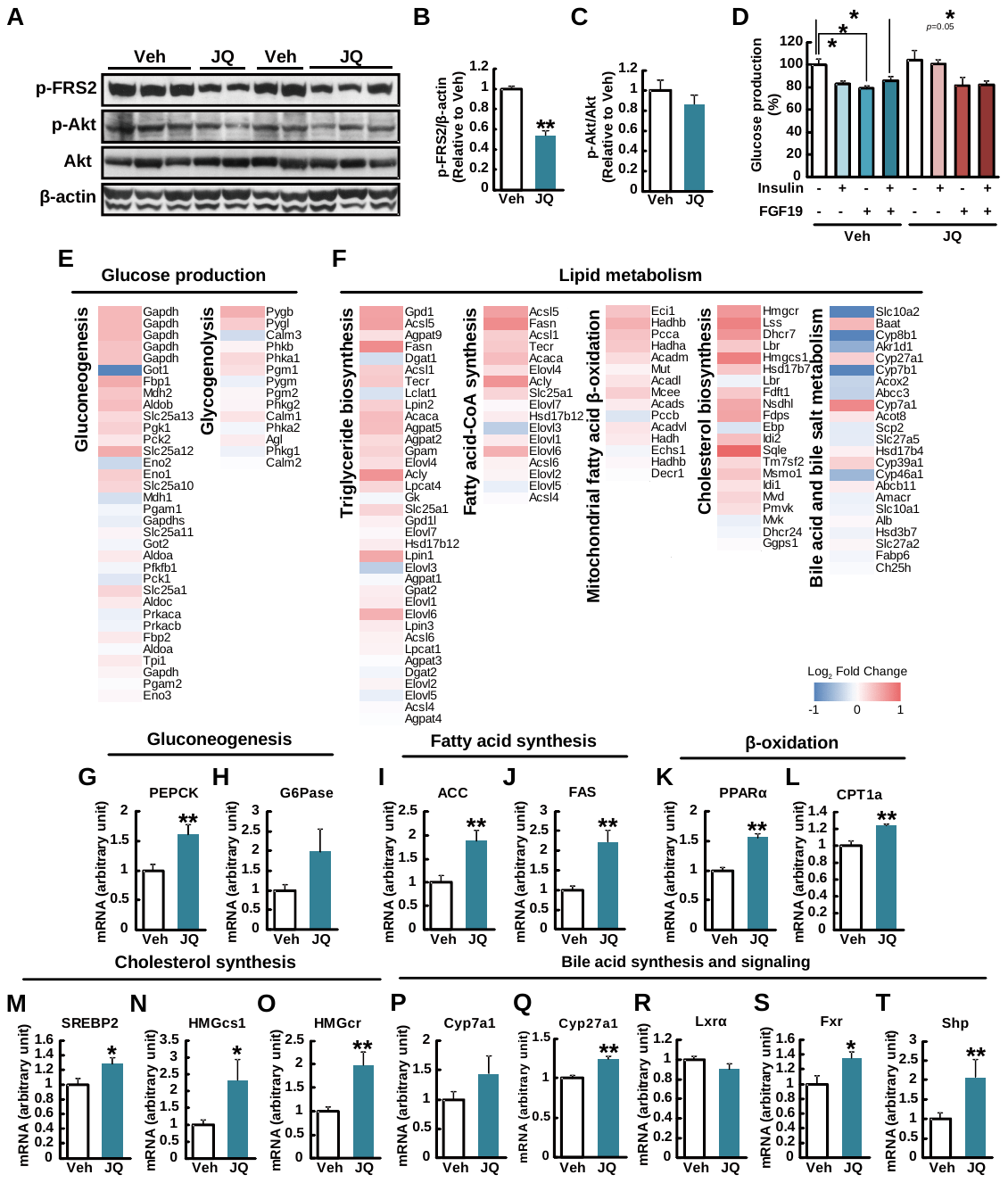

(**A-C**) Western blot analysis of phosphorylated FRS2 (**A, B**) and Akt (**A,** **C**) in liver lysates. β-actin was used as a loading control. (**D**) Glucose production with or without insulin and FGF19 in primary hepatocytes (*n* = 3). **P* < 0.05, ***P* < 0.01, *vs.* vehicle (Veh) or JQ-1-treated cells without insulin or FGF19. Microarray analysis in the liver was performed in JQ-1-treated mice (*n* = 3). Heat maps represent log_2_ fold changes in gene expression in pathways related to (**E**) glucose production, (**F**) lipid metabolism. Expression levels of genes related to (**G, H**) gluconeogenesis (*Pepck* (**G**) and *G6Pase* (**H**)), (**I, J**) fatty acid synthesis (*Acc* (**I**), and *Fasn* (**J**)), (**K, L**) β-oxidation (*Ppara* (**K**), and *Cpt1a* (**L**)), (**M-O**) cholesterol synthesis (*Srebp2* (**M**), *Hmgcs1* (**N**) and *Hmgcr* (**O**)) and (**P-T**) bile acid synthesis and signaling (*Cyp7a1* (**P**), *Cyp27a1* (**Q**), *Nr1h3* (LXRα) (**R**), *Nr1h4* (FXR) (**S**), and *Nr0b2* (SHP) (**T**)) in the liver (*n* = 5-6). Expression data were normalized by those of *Rpl13*. **P* < 0.05, ***P* < 0.01, *vs.* vehicle-treated mice (Veh). Data are expressed as means ± SEM.

**Supplemental Figure 2. JQ-1 modulates hepatic sterol metabolism. *Related to Figure 1.***

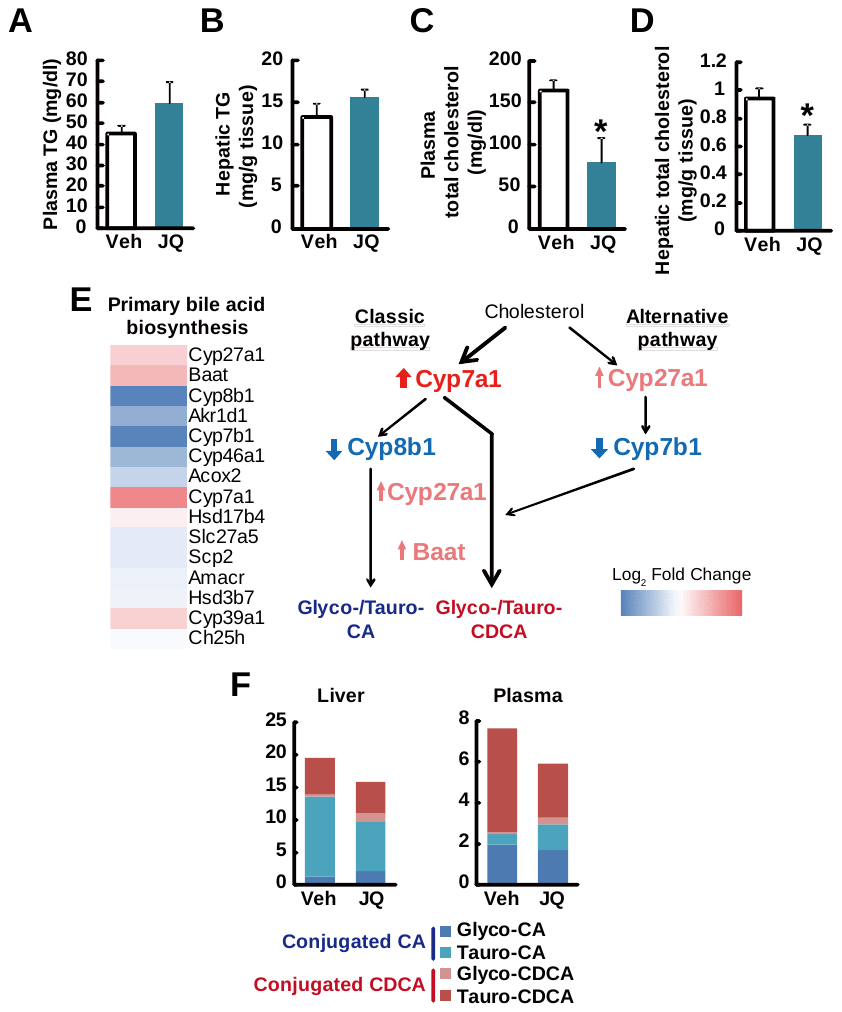

Plasma (**A**, **C**) and hepatic (**B**, **D**) triglyceride (TG) (**A**, **B**) and total cholesterol (**C**, **D**) levels in JQ-1-treated mice (*n* = 5-6). **P* < 0.05, ***P* < 0.01, *vs.* vehicle-treated mice (Veh). (**E**) Heatmap represent log_2_ fold changes in gene expression in “primary bile acid biosynthesis” pathway. (**F, G**) Bile acid composition in the liver (**F**) and plasma (**G**). Data are expressed as means ± SEM.

**Supplemental Figure 3. JQ-1 decreases Reg4+ Paneth cells in the ileum. *Related to Figure 2.***

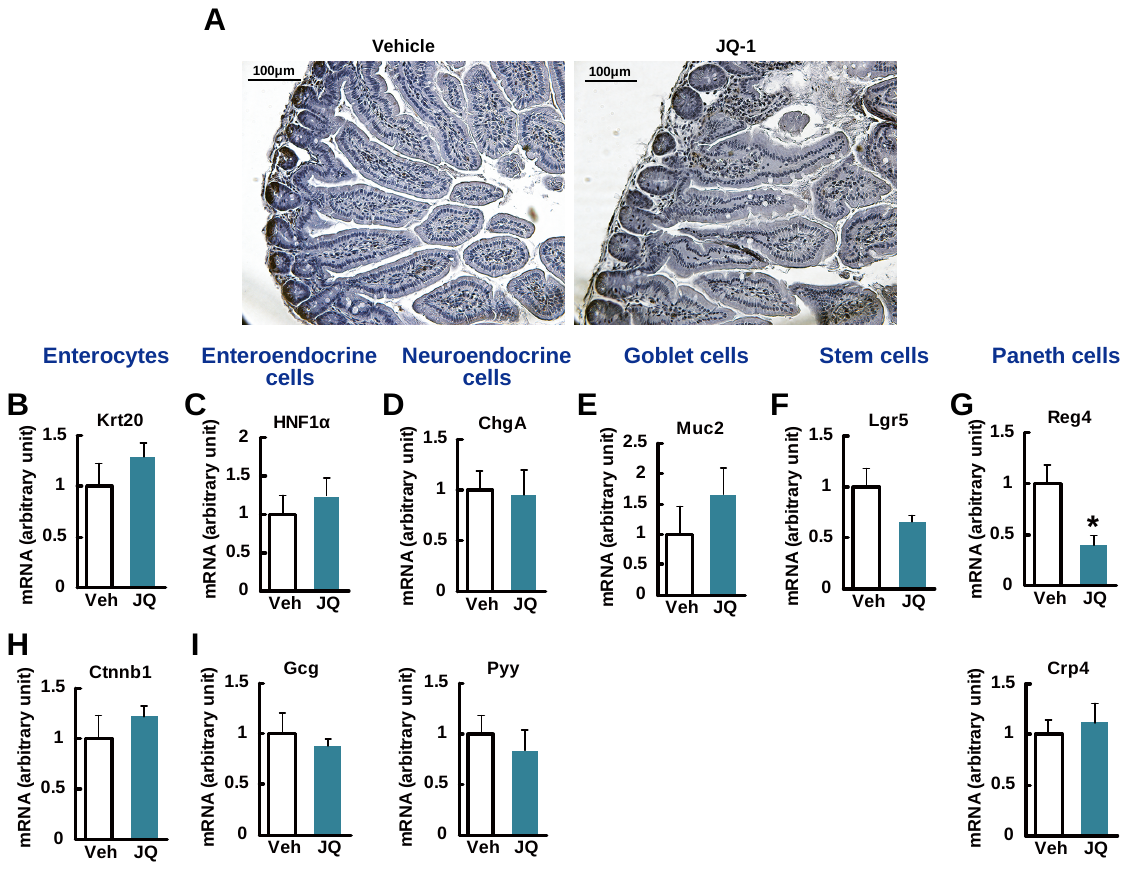

(**A**) Lysozyme staining of intestinal sections. Scale bar, 100 μm. (**B-K**) Gene expression levels of cell type marker genes. Krt20 (**B**) and Ctnnb1 (**H**) are enterocyte markers, HNF1α (**C**) is an enteroendocrine cell marker, and Gcg a secretory product (**I**). ChgA (**D**) is a neuroendocrine cell marker, with Pyy a secretory product (**J**). Muc2 (**E**) and Lgr5 (**F**) are markers of goblet cell and stem cell, respectively. Reg4 (**G**) is a Paneth cell marker, and the cell secretes Crp4 (**K**). **P* < 0.05 *vs.* vehicle (Veh)-treated cells. Data are expressed as means ± SEM.
