## Supplementary material for "Bromodomain inhibition reveals FGF15/19 as a target of epigenetic regulation and metabolic control": Brief Highlights

- Bromodomain inhibition by JQ-1 induces hyperglycemia.
- JQ1 modulates the gut-liver FXR-FGF15/19 signaling pathway.
- JQ-1 reduces Brd4 binding and expression of FGF15 and SHP in ileum.
- JQ-1 effects to induce glucose intolerance are abolished by FGF19 overexpression.
